# A combined program of induced stemness and differentiation in response to IFNγ drives acute myeloid leukaemia growth

**DOI:** 10.64898/2026.09.09.750099

**Authors:** Constandina Pospori, Alessandro Donada, Helena Boutzen, William Grey, Nick Rabas, Sara Gonzalez Anton, Nada Kassara, Wenjie Sun, Flora Birch, Christiana Georgiou, Shayin Gibson, Audrey Au Yong, Adam Karoutas, Tatiana Rizou, Dane Vassiliadis, Georgia Stevens, Thomas Williams, Reema Khorshed, Myriam Haltalli, Alex Murison, Maria-Nefeli Skoufou-Papoutsaki, Katherine Sloan, Hector Huerga Encabo, Jack Hopkins, Chrysi Christodoulidou, Irene Rodriguez-Hernandez, Probir Chakravarty, Dimitris Stampoulis, Samantha Atkinson, Francesca Hearn-Yeates, Aikaterini Polyzou, Eirini Trompouki, Richard Burt, Andrew J. Innes, Hans J. Stauss, Ronjon Chakraverty, Stephanie Z. Xie, Mark Dawson, John E. Dick, Ilaria Malanchi, Leïla Perié, Dominique Bonnet, Cristina Lo Celso

## Abstract

How inflammation shapes acute myeloid leukaemia (AML) has come under scrutiny, as it may explain the disease’s resistance to immunotherapy approaches. IFNγ has emerged as a key cytokine with paradoxical roles in suppressing and supporting AML growth, and the fundamental question of how leukemic stem cells (LSCs) respond to IFNγ and whether IFNγ signaling influences LSC quiescence and their capacity to regenerate disease over the long term remains unanswered. Here, we study primary human AML cells and murine models and combine bioinformatics analyses and functional assays to show that AML hierarchical heterogeneity is responsive to IFNγ challenge. We uncover that IFNγ triggers parallel stemness and differentiation programs; HSC/MPP-like cells enter deeper stemness, associated with quiescence and high leukaemia regeneration potential, while the surviving pool of progenitor-like cells divides faster but produces progeny that is quickly lost. Finally, with murine models we show that exposure to inflammation in vivo results in only transient impairment of leukaemia propagating capacity. This mechanism is driven by a previously unrecognised intraclonal fate bifurcation, relevant for the development of more effective therapeutic approaches.

## Introduction

Acute Myeloid Leukaemia (AML) is an aggressive haematological malignancy with overall consistently poor clinical outcome^1^. While intensive chemotherapy coupled to bone haematopoietic stem cell transplant (HSCT) can be curative, the majority of patients are, elderly and less fit and therefore ineligible for these treatments^1^. Recently developed targeted therapies can extend survival beyond 3-6 months, but are often indicated only for specific molecularly defined AML subsets, and all are marred by resistance^2^. As a result, 5-year survival remains as low as 30% due to incurable relapse^1,2^. AML is composed of hierarchically heterogeneous myeloid leukaemic cells^3–5^, with Leukaemic Stem Cells (LSCs) positioned at the top of the hierarchy, initiating and sustaining disease^6,7^ and driving relapse^8–10^ following therapeutic interventions. Similar to healthy HSCs^11^, more quiescent LSCs exhibit higher self-renewal and long-term repopulating capacity^12^.

Work by us and others has demonstrated that AML induces focal and progressive changes in the Bone Marrow (BM)^13–15^ and therefore grows in an evolving tumour microenvironment. In parallel with BM stroma and haematopoiesis remodelling^13–15^, evidence of anti-tumour T cell and NK cell responses have been reported both in patients and murine models of AML^16–22^. These immune responses are eventually unsuccessful in controlling disease, and T cell infiltration, IFNγ signalling and IFNγ-driven transcriptional signatures have been emerging as prognostic indicators of resistance to venetoclax^23^ and chemotherapy^24,25^, but also of responsiveness to immunotherapy^24,26^. Both IFNγ administration and inhibition have been proposed as AML therapeutics^23,25,27^, and opposite effects have been linked to IFNγ dose^28^. This emerging complex picture highlights the unmet need to better understand this cytokine’s multifaceted effects on AML cells. Several studies demonstrate the impact of inflammation and particularly IFNγ on healthy haematopoiesis^29–36^, and inflammation-resistant HSCs have been identified in human and mice as drivers of regeneration^11,35,38^. These studies raise the fundamental question of how leukemic stem cells (LSCs) respond to IFNγ and whether IFNγ signaling influences LSC quiescence and their capacity to regenerate disease over the long term. We interrogated human primary cells using a combination of single cell bioinformatics and functional approaches. We uncovered that acute IFNγ challenge triggers parallel stemness and differentiation programs within clonally related AML cells in primary samples. Resulting LSCs become deeply quiescent and exhibit transcriptional signatures of strong leukemogenic function, while differentiation is linked to both clonogenic cell proliferation and cell loss among their progeny. Murine models demonstrate the role of these mechanisms *in vivo*, whereby IFNγ drives initial reduction of primitive cells compartment size and function, followed by eventual rebound and LSC expansion. These findings explain how the initial reduction in disease growth is followed by eventual rebound and reconcile the contrasting roles of IFNγ reported in literature.

## Results

### IFNγ triggers parallel stemness and differentiation programs in CD34+ human primary AML cells

We selected a number of primary patient samples covering a range of cytogenetic subtypes (Table 1). First, we exposed bulk primary AML samples (n=6) to IFNγ/PBS in culture for three days and confirmed MHC II and PDL1 upregulation in both CD34^+^ (known to be enriched in AML stem and progenitor cells^6,7^) and CD34-sub-populations (Supplementary Figure 1a-d). This suggested that both sub-populations responded to the inflammatory stimulus. CD38 upregulation resulted in a statistically significant decrease in the proportion of CD34^+^ CD38^−^ AML cells (Supplementary Figure 1e). To gain a deep understanding of the molecular processes triggered by IFNγ in LSC-enriched AML cells we performed single nuclei 10x Multiome sequencing (snRNA-seq + snATAC-seq) on CD34+ cells from 4 patient samples, FACS-purified from our IFNγ/PBS bulk cultures. Following batch correction between patient samples, all data were integrated and cells from each patient and condition were mapped onto the human haematopoiesis, single-cell transcriptional atlas^5^ (Figure 1a). This allowed us to identify multiple populations within our samples, starting from HSC-like, multipotent progenitors (MPP) and lympho-myeloid primed progenitors (LMPP)-like and then progenitor and differentiated-like cells bifurcating either towards a megakaryocyte/erythroid differentiation trajectory (MEP, EoBasoMast Precursor, Megakaryocyte precursor, early erythroid and late erythroid cells) or a monocyte differentiation trajectory (cycling progenitors, early GMP, late GMP, pro-Monocyte, monocyte and cDC) (Figure 1a). Each AML displayed distinct cellular hierarchy patterns (Figure 1b, left-side column PBS condition). Interestingly, cells exposed to IFNγ appeared to accumulate towards the tips of their differentiation landscape, mapping either over HSC/MPP cell states or over more differentiated cell states (Figure 1b, middle column). To compare the distribution of PBS and IFNγ treated CD34^+^ AML cells across the differentiation trajectory irrespective of lineage output, we calculated a pseudotime score predicting how advanced in the reference map trajectory cells were, for each sample in the two conditions (Figure 1b – right side panels). Cells on the left side, with the lowest pseudotime scores, were closest to the most primitive HSC population transcriptional profile (HSC/MPP-like cells); cells on the right side, with the highest scores, were farthest from the HSC population, and thus the most mature (differentiated-like cells); cells in the middle would have intermediate scores (progenitor-like). Indeed, we observed a statistically significant enrichment in HSC/MPP-like cell states among IFNγ treated CD34+ AML cells compared to PBS ones (Figure 1c), coupled with a trend for fewer cells in progenitor-like states (Figure 1d) and more cells in differentiated-like cell states (Figure 1e). Of note, progenitor-like cells were the bulk of the overall CD34+ population in both PBS and IFNγ conditions. This analysis highlighted a pattern among CD34+ leukaemic cells to be residing at the two extremes of their differentiation trajectories following IFNγ treatment.

**Figure 1.**
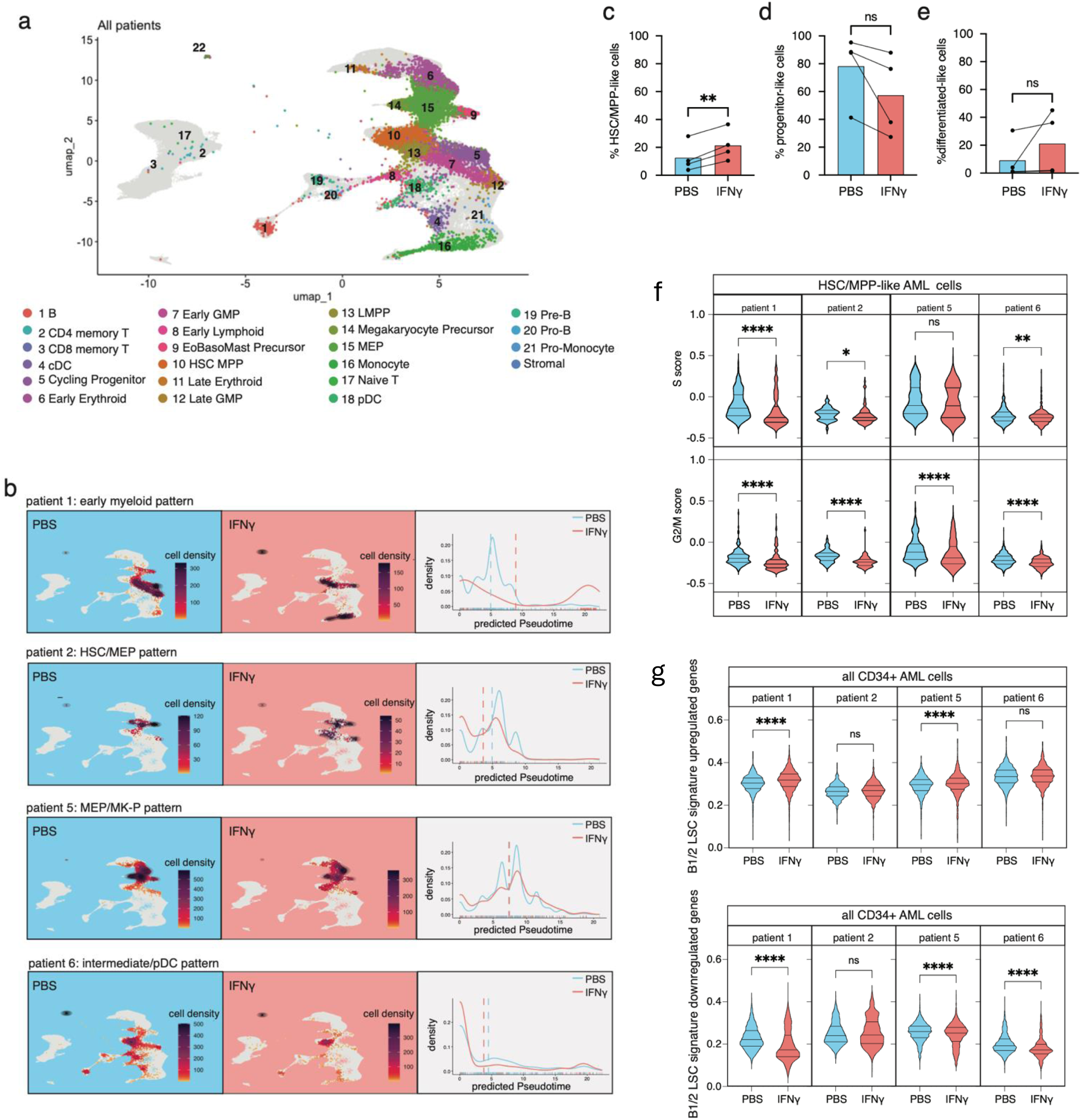
**a,** snRNAseq data of FACS sorted CD34^+^ AML cells from 4 primary AML patient samples treated with PBS or 50ng/mlIFNγ for 3 days *in vitro*, were mapped onto the BoneMarrowMap^5^ as described in the method section. Cells were coloured based on the closest cell type they mapped onto the BoneMarrow reference Map, independently of the patient sample they are coming from. Cells part of the reference map is colored in grey in the background to visualise where AML patient cells land **b,** For each patient, PBS-treated (left-side panels, blue boxes) and IFNγ-treated (middle panels, coral red boxes) CD34^+^ cells mapped to the BoneMarrowMap^5^ are shown as coloured dots, with colour indicating cell density. Right side panels depict the pseudotime trajectory analysis of the same data (PBS:blue line, IFNγ: red line). **c,** Percentage of cells mapped on HSC/MPP-like cell states among CD34^+^ AML cells sort purified and sequenced from PBS and IFNγ treated patient samples. **d,** Percentage of cells mapped on progenitor-like cell states among CD34^+^ AML cells sorted and sequenced from PBS and IFNγ treated patient samples. **e,** Percentage of cells mapped on differentiated-like cell states among CD34^+^ AML cells sorted and sequenced from PBS and IFNγ treated patient samples. in (c-e) each dot represents a different patient and paired t-tests performed for statistical significance. **f,** S-phase (top row) and G2/M (bottom row) signature scores on HSC/MPP-like cell state cells among sorted and snRNA sequenced CD34^+^ AML cells from each of the 4 patient samples exposed to 50ng/ml rhIFNγ or PBS control. t-tests with Mann Whitney correction performed for statistical significance. **g,** B1/2 LSC upregulated genes signature scores (left side-panel) and downregulated signature scores (right side-panel) on all FACS sorted and scRNA sequenced CD34^+^ AML cells from each of the 4 patient samples exposed to 50ng/ml rhIFNγ or PBS control. t-tests with Mann Whitney correction performed for statistical significance.

**Table 1.**
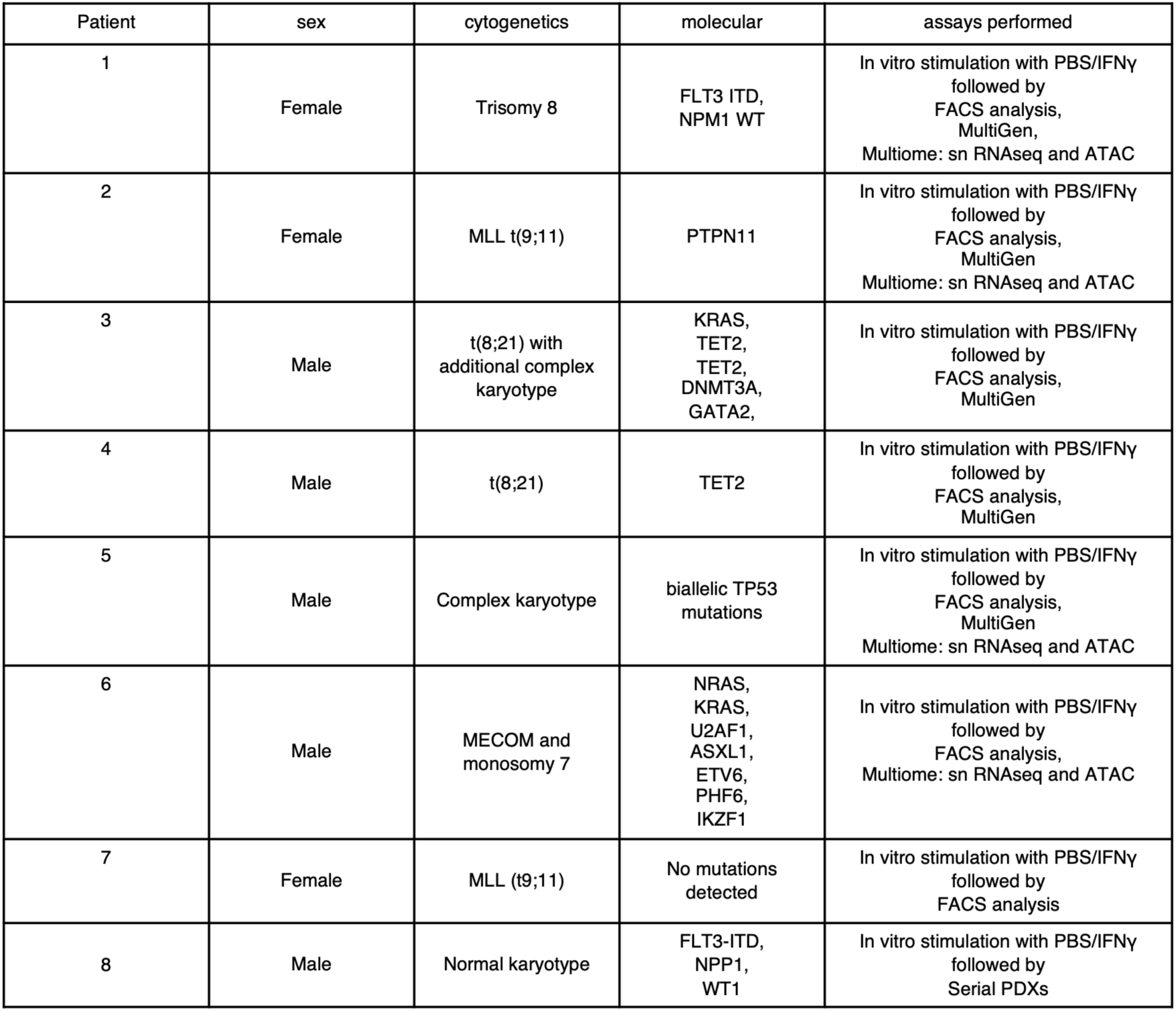
Patient sample details.

| Patient | sex | cytogenetics | molecular | assays performed |
|---|---|---|---|---|
| 1 | Female | Trisomy 8 | FLT3 ITD,<br>NPM1 WT | In vitro stimulation with PBS/IFNγ<br>followed by<br>FACS analysis,<br>MultiGen,<br>Multiome: sn RNAseq and ATAC |
| 2 | Female | MLL t(9;11) | PTPN11 | In vitro stimulation with PBS/IFNγ<br>followed by<br>FACS analysis,<br>MultiGen<br>Multiome: sn RNAseq and ATAC |
| 3 | Male | t(8;21) with additional complex karyotype | KRAS,<br>TET2,<br>TET2,<br>DNMT3A,<br>GATA2, | In vitro stimulation with PBS/IFNγ<br>followed by<br>FACS analysis,<br>MultiGen |
| 4 | Male | t(8;21) | TET2 | In vitro stimulation with PBS/IFNγ<br>followed by<br>FACS analysis,<br>MultiGen |
| 5 | Male | Complex karyotype | biallelic TP53 mutations | In vitro stimulation with PBS/IFNγ<br>followed by<br>FACS analysis,<br>MultiGen<br>Multiome: sn RNAseq and ATAC |
| 6 | Male | MECOM and monosomy 7 | NRAS,<br>KRAS,<br>U2AF1,<br>ASXL1,<br>ETV6,<br>PHF6,<br>IKZF1 | In vitro stimulation with PBS/IFNγ<br>followed by<br>FACS analysis,<br>Multiome: sn RNAseq and ATAC |
| 7 | Female | MLL (t9;11) | No mutations detected | In vitro stimulation with PBS/IFNγ<br>followed by<br>FACS analysis |
| 8 | Male | Normal karyotype | FLT3-ITD,<br>NPP1,<br>WT1 | In vitro stimulation with PBS/IFNγ<br>followed by<br>Serial PDXs |

We previously showed that among functionally heterogeneous LSC populations^12,39^, the LSCs exhibiting superior long-term repopulating activity are quiescent, and when examined in clonogenic assays they produce smaller, less differentiated progeny than more proliferative LSCs which initially reconstitute xenografts faster but are eventually outcompeted by quiescent LSCs^12^. Taking this work into consideration, together with the fact that IFNγ has been documented to have cytostatic effects in multiple cancers^40,41^ including AML^25^, we asked whether HSC/MPP-like cells are not only numerically enriched among sorted CD34+ AML cells post-IFNγ but also become more quiescent. Indeed, when comparing cell cycle signature scores between the PBS and IFNγ-treated cells assigned to the HSC/MPP-like cell state, the latter were shown to express lower S and G2/M phase signature scores (Figure 1f).

To confirm the link between deep quiescence and stemness in LSCs that survived IFNγ exposure, we turned to gene signatures generated through the comparison of two functionally validated AML LSC subsets identified in the OCI-AML22 human AML model^12,42^; one subset being deeply quiescent, with enhanced self-renewal, better long-term leukaemia repopulating capacity and chemoresistance, and the other more proliferative and producing more differentiated progeny. The B1/2 LSC signature containing the list of genes upregulated in the deeply quiescent and chemoresistant LSC subset^12^ were higher in the CD34^+^ AML cells from IFNγ-treated patient samples and this was statistically significant in 2 out of the 4 samples examined (Figure 1g left side panel). The B1/2 LSC signature containing the list of downregulated genes in this population was lower in the CD34^+^ AML cells from the IFNγ-treated samples compared to the PBS ones, and this was statistically significant in 3 out of 4 samples (Figure 1g, right side panel). This analysis points to a mechanism through which some LSCs may survive acute IFNγ inflammation by entering a deeper state of quiescence and self-renewal, and raises the question whether this may be achieved through different LSC-containing cell families undergoing differentiation while stemness is maintained or triggered in others.

### AML clones surviving inflammatory insult contain leukaemia stem cells with high functional potential

The differentiation trajectory redistribution patterns we observed at the population level (Figure 1b-e) can be the result of isogenic cells responding to IFNγ in different ways with some LSCs induced to enter deeper stemness states, and progenitor-like cells induced to either die or differentiate. Alternatively, the IFNγ-induced enrichment in cell states at the two ends of the differentiation trajectory might be the result of distinct clonal families responding differently to IFNγ. To explore which mechanism is more likely, we used mitochondrial DNA mutations as natural barcodes, to identify cells that are clonally related^43^. For each of the three patient samples displaying sufficient mitochondrial mutations for lineage tracing, we identified cells that had a high probability to be clonally related based on mtDNA mutation clustering analysis. Three clonally related cell clusters were identified in patient sample 1, two in patient sample 5 and four in patient sample 6, as shown in heatmaps of binarized mutation status for each mitochondrial locus in each cell within each patient (Supplementary Figure 2b), clustered to produce heatmaps of binarized mutation status in cells belonging to clonally-related clusters within each patient (Figure 2a). Next, we mapped PBS and IFNγ-treated cells belonging to each clonally-related cell cluster on the predicted pseudotime trajectory (Figure 2a histograms). Broadly, individual clonally-related cell clusters appeared to distribute across the predicted pseudotime similarly to how all PBS/IFNγ cells from the corresponding patient samples did (Figure 2a histograms and Figure 1b right panels). This was also the case, when we examined cells grouped together based on commonly expressing each individual mitochondrial mutation (Supplementary Figure 2c). Interestingly, all individual, clonally-related cell clusters identified in AML samples 1 and 6, which more strongly polarised towards the edges of the differentiation trajectory and predicted pseudotime in the IFNγ condition (Figure 1b), contained cells at the HSC/MPP-like edge of the predicted pseudotime, but only some contained cells at the other end, corresponding to differentiated-like cells (Figure 2a, patient 1 and patient 6 samples). Overall these data indicate that differentiation at the population level was not driven by some distinct, differentiated clones, but rather was a component of observed clones and always coupled with stemness.

**Figure 2.**
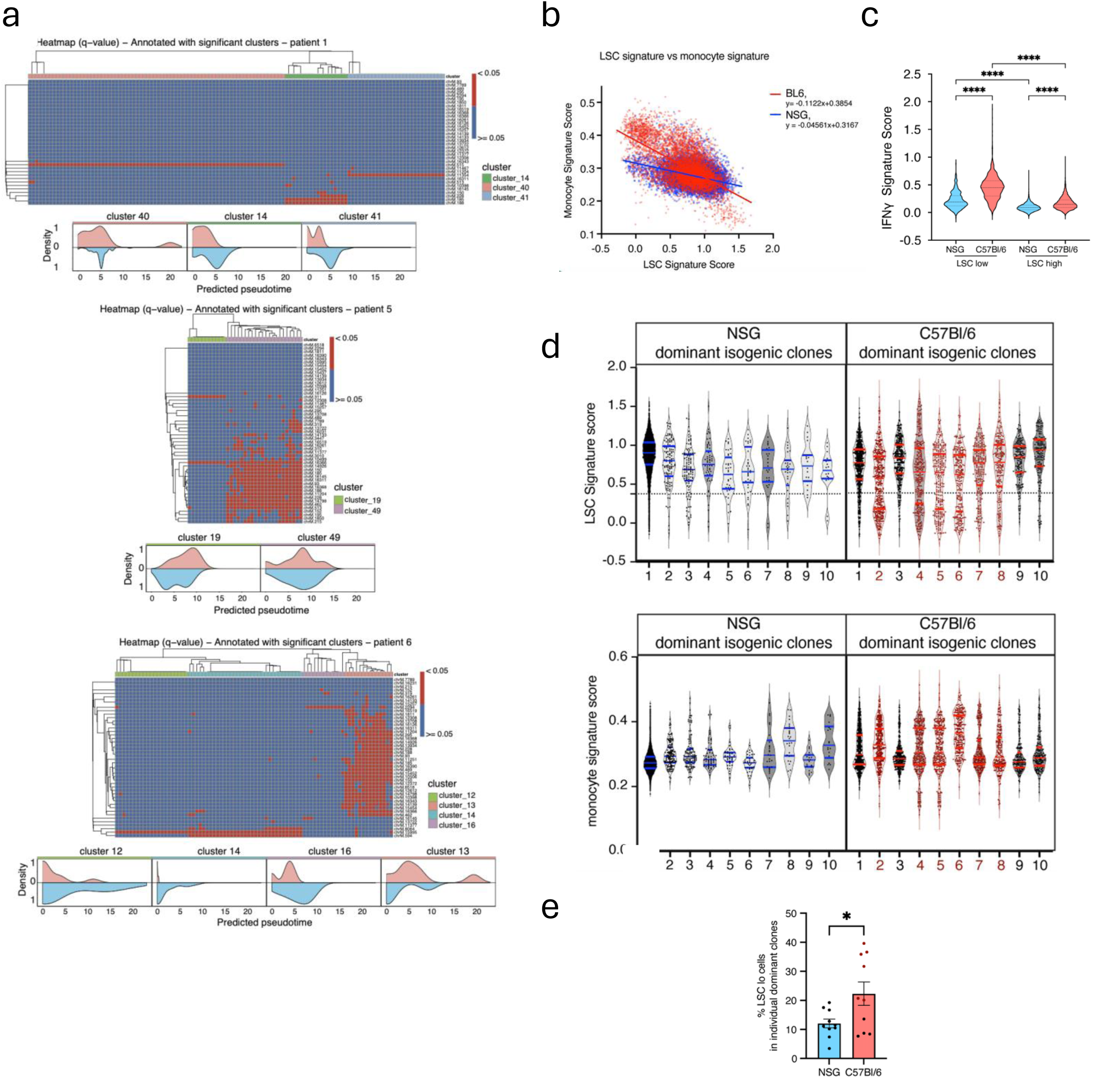
**a,** Heatmap showing the binarised mutation status for each mitochondrial locus in each cell, displayed per patient, and annotated by clonally-related cell cluster. Columns represent cells and rows represent mitochondrial loci. Loci with q-value < 0.05 are shown in red (mutated), while loci with q-value ≥ 0.05 are shown in blue (non-mutated or uncertain). Cells were clustered using hierarchical clustering with Jaccard distance and average linkage. Among the clusters with strong bootstrap support (AU p-value ≥ 0.95), to identify clonally-related cell clusters within each patient. Only clonally-related cell clusters containing at least 10 cells in total and at least 2 cells per condition are displayed. Below the heatmap from each of the 3 patient samples, density plots show the distribution of cells for each identified clonally-related cell cluster along the predicted pseudotime, for condition IFNγ in red, and PBS in blue. **b,** Correlation between LSC signature score and monocyte signature score of *MLL-AF9 + Kras^G^*^12^ AML cells from NSG (blue dots) and C57Bl/6 (red dots) leukaemic animals. Simple linear regression test performed for statistical significance. **c,** IFNγ response signature scores on LSC^high^ and LSC^low^ AML cells from NSG and C57Bl/6 recipients. Dunn’s multiple comparisons and Kruskal-Wallis tests performed for statistical significance. **d,** LSC and monocyte signature scores (top and bottom row panels respectively) of the top 10 individual dominant clones of *MLL-AF9 + Kras^G^*^12^ AML spleen cells from NSG (left side panels, blue) and C57Bl/6 recipients recipients (right side panels, red). Dots within each clone represent individual cells. Quartiles (thick lines) and median (thin red lines) values shown for each clone (blue in NSG, red in C57Bl/6). Dotted line denotes the cut-off for categorising cells as LSC^low^/LSC^high^ (set at the value of 0.375). **e,** Percentage of LSC^low^ signature score cells within each of the top 10 dominant clones in NSG (blue) and C57Bl/6 recipients (red). Dots are individual clones, error bars: mean ± SEM. C57Bl/6 dominant clones containing more than 20% LSC^low^ cells shown in maroon.

To extend our human AML computational findings into a system more tractable for clonal interrogation, we turned to an existing dataset of barcoded murine AML where cell clones were easily distinguishable. MLL-AF9-Kras*^G12D^*AML cells were modified to express SPLINTR barcodes, expanded *in vitro* and transplanted into NSG and C57Bl/6 recipients to model AML growth in inflammation-free and inflammatory conditions *in vivo*, respectively. While different clones expanded in the the two types of recipients, these were otherwise isogenic^44^. First, we evaluated the LSC, monocyte differentiation and IFNγ differentiation score of all cells harvested from fully infiltrated NSG and C56Bl/6 recipients. MLL-AF9-Kras*^G12D^* AML cells harvested from fully infiltrated C57Bl/6 recipients had an overall lower LSC signature score and a higher monocyte signature score than the same cells grown in NSG recipients and the two scores were inversely correlated in both groups, though this was more prominent for cells growing in C57Bl/6 animals (Figure 2b). Importantly, both LSC high and LSC low scored cells from C57Bl/6 mice exhibited higher IFNγ response signature than their counterpart from NSG recipients (Figure 2c), consistent with IFNγ playing a role in the transcriptional changes observed. Again, differences in LSC and monocyte scores at the population level could be driven either by the coexistence of stem cell clones and differentiated clones of varying sizes, or by reactive remodelling of the hierarchical heterogeneity of each individual clone across the population. To investigate this, we assessed these signatures in individual SPLINTR-barcoded, dominant isogenic clones of MLL-AF9-Kras*^G12D^* AML cells harvested from NSG or C57Bl/6 recipients^44^. Most cells in each of the top 10 dominant clones grown in NSG mice exhibited high LSC signature and low monocyte signature scores (Figure 2d, left side panels, top and bottom rows), but C57Bl/6 dominant clones were more diverse (Figure 2d, right side panels, top and bottom rows). 6 out of 10 clones from C57Bl/6 recipients exhibited a bimodal distribution across LSC and monocyte signature scores with more than 20% of cells expressing an LSC^low^ signature score (Figure 2e), indicating a cellular hierarchy, heavier at its two extremes, linked to the C57Bl/6 microenvironment. Importantly, all dominant C57Bl/6 clones harvested at full infiltration contained an LSC^high^ cell population (Figure 2d, top row, right side panel).

Both human and murine analyses identified that no clonally related cell clusters that survived IFNγ exposure where solely enriched in differentiated cells. This suggested that individual AML clones surviving the inflammatory insult are poised to remain long-term, which would require testing using functional assays.

### IFNγ acutely reduces LSC clonogenic capacity through loss of progeny and delayed stem cell expansion

To examine the functional impact of acute IFNγ on human CD34^+^ AML cells at the single cell level, we treated 5 AML samples (Table 1) with PBS/IFNγ for 3 days, then assessed single, FACS purified CD34^+^ cells with the MultiGen single cell fate mapping assay (Supplementary Figure 2), which uniquely combines the immunophenotypic characterisation of the progeny of individual clonogenic cells with a record of their division history within a 72hr window^45–47^ and therefore analyses cell cycle at the functional level. The number of clonal families recovered from CD34^+^ primary AML cells exposed to IFNγ (‘IFNγ families’ from now on) was reduced (Figure 3a-b), IFNγ families showed a trend towards being smaller (Figure 3a and 3c), and the overall number of AML cells recovered per IFNγ-exposed cell plated was markedly reduced in each patient (Figure 3d). Surprisingly, division history analysis revealed that in 4 out of 5 patient samples a higher percentage of cells in IFNγ families progressed to later generations within the 72hr window (Figure 3e), with a higher Mean Division Number (MDN) score confirming faster division (Figure 3f). A possible explanation for faster cell division not resulting in a greater clonal output could be increased cell death. As a proxy measure for cell death, which is not directly assessed in the MultiGen assay, we examined individual PBS/IFNγ family completeness, defined as the percentage of cells recovered out of the number of cells expected per clonal family based on the number of divisions measured. IFNγ-families were less complete than PBS ones (Figure 3g), especially in later generations (3 and above; Figure 3h), consistent with the hypothesis that while CD34^+^ AML cells which continued to cycle immediately after IFNγ exposure divided faster, later generation (more rapidly dividing) cells were more susceptible to cell death. Finally, the percentage of CD34^+^ progeny per clonal family was not significantly different between the PBS and IFNγ pre-treatment conditions at the end of the assay (Figure 3i), and this was also the case at the population level (Figure 3j). Overall, our findings suggested that acute exposure to IFNγ markedly reduces CD34^+^ AML growth by impairing both clonogenicity and progeny viability, but without altering rate of exit from the CD34^+^ compartment.

**Figure 3.**
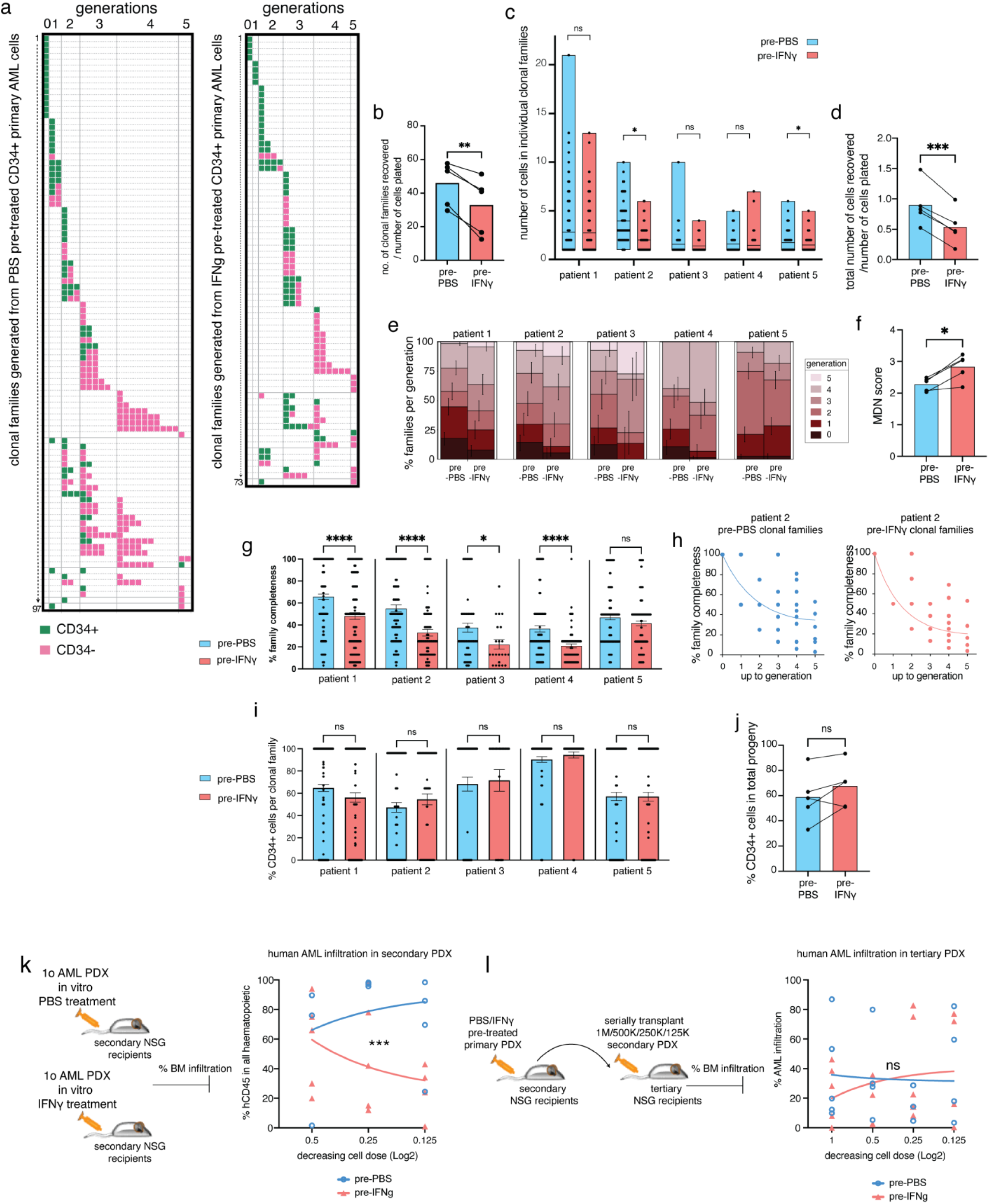
**a,** Representative (patient 2) division history and phenotype of individual progeny of PBS (left-side panel) and IFNγ pre-treated (right-side panel) CD34^+^ primary AML cells. Each row represents a clonal family, each square represents a single cell. Green squares: CD34^+^, pink squares: CD34^−^. **b,** Percentage of clonal families recovered per number of PBS/IFNγ pre-treated CD34^+^ cells plated. n=5 independent primary AML samples. Paired t-test performed for statistical significance. **c,** Number of cells in individual clonal families. Dots indicate individual clonal families, lines at mean. t-tests with Welch correction performed for statistical significance. **d,** Total number of cells recovered. Dots represent each patient sample examined, paired t-test performed for statistical significance. **e,** Percentage of clonal families containing cells of each generation. Error bars: mean +/− SEM. Permutation-based test to challenge the null hypothesis that the distribution of clonal families across generations was independent of IFNγ exposure was performed for statistical significance, (patient 1: p-value = 0.012, n=313 clonal families, patient 2: p-value = 0.086, n=170 clonal families, patient 3: p-value=0.005, n=78 clonal families, patient 4: p=0.017, n= 165 clonal families, patient 5: p-value=0.039, n=302 clonal families) **f,** MDN score for the clonal progeny of PBS/IFNγ pre-treated CD34^+^ cells in each patient sample. Dots represent individual patient samples, paired t-test performed for statistical significance. **g,** Percentage family completeness, calculated as the number of cells in individual clonal families divided by the number of cells expected in a complete clonal family reaching the same generation as the one examined. Error bars: mean +/− SEM, t-tests with Welch correction performed for statistical significance. **h,** Percentage clonal family completeness in relation to the generation reached by each clonal family recovered from PBS (left-side panel) /IFNγ (right-side panel) pre-treated CD34^+^ plated cells, in a representative patient sample. Each dot represents an individual clonal family, line: non-linear regression model. **i,** Percentage of CD34^+^ cells in each clonal family. Error bars: mean +/− SEM, t-tests with Welch correction performed for statistical significance. **j,** Percentage of CD34^+^ cells in each patient sample examined. Dots represent individual patients. paired t-test performed for statistical significance. **k,** BM cells from a primary PDX sample were cultured for 3 days in the presence of IFNγ or PBS control and subsequently transplanted into NSG recipients at three doses. Human AML BM infiltration was assessed at 11 weeks post-transplant and human CD45^+^ cells engraftment is reported. 3 cell doses used: 500,000 per mouse, n=5 per group, 2 mice receiving PBS-pretreated cells were fully infiltrated and had to be sacrificed at an earlier time point, 250,000 cells per mouse, n=5 per group and 125,000 cells per mouse, n=4 mice receiving PBS-pretreated cells and n=5 mice receiving IFNγ-pretreated cells. **l,** Serial transplantation of PBS/IFNγ pre-treated human AML cells. Percentage of human CD45^+^ cells among all haematopoietic cells (mouse+human) in each recipient shown in adjacent graph. k and l, Each dot represents a recipient animal. Simple linear regression shown. ANCOVA (analysis of covariance) test performed for statistical significance.

LSCs - defined as cells with *in vivo* leukaemia initiating capacity in mouse xenografts - are known to be rare even within the CD34^+^ compartment. Because the MultiGen assay fate maps most efficiently cells that proliferate within a short-term period (72hr), its results describe the impact of IFNγ on the output of more proliferative, progenitor-like AML cells. To better assess whether human AML LSCs can recover from IFNγ exposure, we established a primary patient derived xenograft (PDX) (patient sample 1, table 1), harvested fully infiltrated BM, exposed the leukaemic cells to IFNγ or PBS *in vitro* for 3 days, and then re-injected them into secondary NSG recipients (Figure 3k). 11 weeks later, AML infiltration was lower in recipients of IFNγ pre-treated cells (Figure 3k), an indication of reduced repopulating activity. However, AML cells harvested from secondary PBS/IFNγ group recipients grew equally well in tertiary recipients (Figure 3l), indicating that AML LSCs function recovered in the long-term. Combined, the MultiGen and PDX data showed that acute IFNγ exposure causes a reduction in proliferative CD34+ AML cells, that those that survive proliferate faster but their progeny has reduced survival. However some quiescent LSCs survive and are able to sustain serial PDX reconstitution.

### In vivo exposure to IFNγ affects LSC function only temporarily

To investigate whether exposure to IFNγ would functionally affect LSCs *in vivo,* we generated an immunogenic murine AML model. We transduced MLL-AF9 AML cells with the influenza A nucleoprotein (NP) model-antigen (NP-MLL-AF9 AML, from here on NP-AML), which is recogonised by endogenous NP-specific C57Bl/6 T cells^48,49^, and also by all T cells from F5 TCR transgenic C57Bl/6 mice. In co-culture experiments, NP-AML cells activated T cells from F5 TCR transgenic animals but not control OTI transgenics, resulting in a much lower number of AML cells at the end of the culture, and expansion of F5 but not OTI T cells (Supplementary Figure 4a). Interestingly, the AML cells co-cultured with F5 T cells had. Alower proportion of c-Kit+ cells, enriched for LSCs in this model^50,51^ (Supplementary Figure 4b). Conditioned media from the F5:NP-AML co-cultures, but not the OTI:NP-AML co-cultures, led to a reduction in the colony forming capacity of the leukaemic cells, which was restored when IFNγ was blocked (Supplementary Figure 4c), and exposing NP-AML cells directly to IFNγ *in vitro* resulted in PDL1 upregulation and reduction of c-Kit+ AML cells (Supplementary Figure 4d). Finally, when we injected NP-AML cells in C57Bl/6 mice we could detect activation and expansion of the otherwise undetectable population of endogenous NP-specific T cells (Supplementary Figure 4e, left). Importantly, while the remaining T cells showed varying expression of CD44 and PD1, all NP-specific T cells were double positive and had the highest PD1 MFI, confirming their initial activation and eventual exhaustion^52^ (Supplementary Figure 4e, right).

To test the effect of *in vivo* IFNγ exposure on LSCs in the shorter and longer term, we injected NP-AML cells in immunocompetent C57Bl/6 and immunodeficient NSG mice and assessed the phenotype of leukaemic cells harvested from fully infiltrated mice from the two groups (NSG on day 17, C57Bl/6 on day 25; cryopreserved and analysed together; Figure 4a-b). Flow cytometry analysis revealed that, while BM infiltration was approximately 95% in all mice (Figure 4c, left), c-Kit expression strikingly differed in the two groups, with a much higher proportion of AML cells from NSG mice expressing c-Kit (Figure 4c, middle and right panels). Next, we functionally assessed individual, FACS-purified c-Kit^+^ AML cells using single-cell colony forming assays. The colony forming capacity (CFU-C) of cells from immunocompetent animals was reduced (Figure 4d) and this was coupled with a small but significant reduction in the size of individual colonies (Figure 4e), suggesting both LSC frequency and proliferative capacity were impaired following exposure to an inflammatory environment. However, the percentage of c-Kit^+^ AML cells in C57Bl/6-derived colonies matched that observed in colonies from NSG-harvested c-Kit^+^ cells (Figure 4f). Consistent with the enrichment in highly functional LSC signatures observed in human CD34+ AML cells exposed to IFNγ in culture, the secondary colony forming capacity of c-Kit^+^ AML cells from C57Bl/6 derived primary colonies was higher than from NSG derived ones (Figure 4g). This confirmed that the few LSCs that retained clonogenic capacity in the aftermath of inflammatory stress, although immediately less proliferative *ex vivo*, displayed enhanced long term self-renewal by regenerating a prominent and potent LSC compartment.

**Figure 4.**
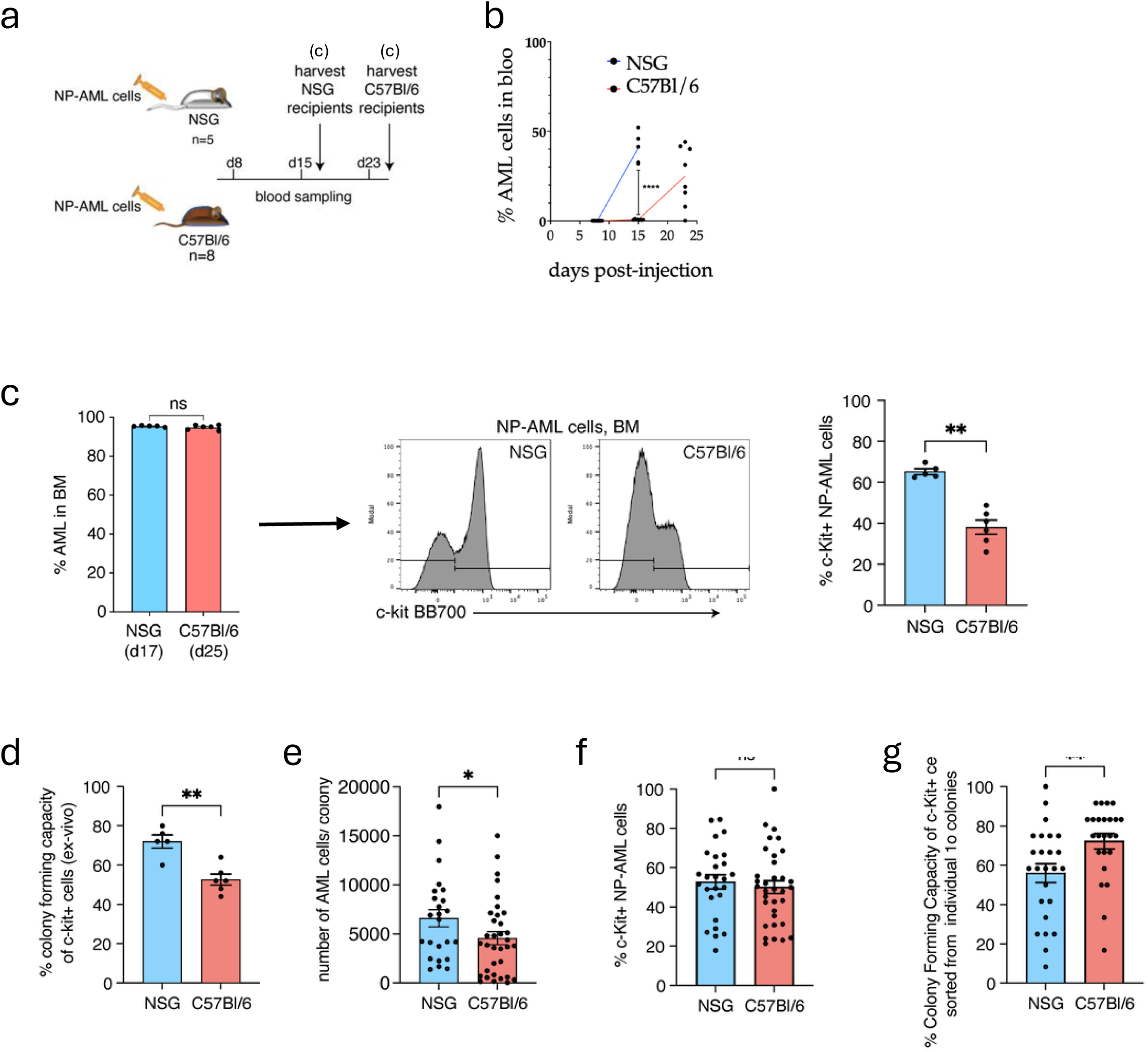
**a-b,** 100,000 NP-AML cells were injected into NSG (n=5) and C57Bl/6 recipients (n=8). Blood AML infiltration was monitored on days 8, 15 and 23. All NSG animals were fully infiltrated and sacrificed on day 17. C57Bl/6 mice were sacrificed on day 25, with 6 out of 8 mice being fully infiltrated at that time point. **c,** BM cells from fully infiltrated animals were cryopreserved and later thawed for immunophenotypic and functional assays. Left: AML infiltration; middle: representative histograms of c-Kit expression; right: percentage of c-Kit+ AML cells from NSG and C56Bl/6 recipients. Dots represent individual mice, error bars: mean ± SEM. Mann-Whitney test performed for statistical significance. **d,** 25 c-Kit+ AML cells from (c) were FACS sorted into single cell, serial colony forming assays. Colony forming capacity of FACS purified c-Kit^+^ AML cells from fully infiltrated NSG and C57Bl/6 leukaemic animals from (d). Dots represent data from individual mice. Error bars: mean ± SEM. Unpaired t-test with Welch correction performed for statistical significance. **e,** Number of cells in individual primary colonies. **f,** percentage of c-Kit^+^ AML cells in individual primary colonies. **g,** secondary colony forming capacity of c-Kit^+^ AML cells sorted from individual primary AML cell colonies. for (e-g) n= 24 primary colonies growing from single c-Kit^+^ AML cells harvested from NSG mice (5 colonies/mouse for 4 animals and 4 colonies/mouse for 1 animal) and n= 34 colonies growing from c-Kit^+^ AML cells harvested from C57Bl/6 mice (6 colonies/mouse for 5 mice and 4 colonies/mouse for 1 mouse). Each dot represents an individual colony. Error bars: mean ± SEM. Mann-Whitney test performed for statistical significance.

## Discussion

IFNγ is widely recognised as a cytostatic agent^40,41^, but also as an inducer of differentiation and stemness loss on healthy HSC populations^29,32–35^. Very recent evidence has been highlighting rare HSCs that retain stemness despite mounting IFN responses^35,38^. The effect of IFNγ on AML has been reported to be both supportive of and detrimental to disease growth^25,28^. Here we reconcile these observations by combining human and murine, *ex vivo* and *in vivo*, population and single cell analyses. snRNA/ATAC multiome sequencing and MultiGen functional analyses of CD34+ primitive cells from cytogenetically distinct AML cases following IFNg challenge revealed two co-existing patterns of behaviour. More abundant, proliferating progenitor cells that survive IFNγ challenge divide more rapidly than vehicle treated cells but give rise to less complete cell families, which remain smaller as a result. CD34^+^ cells upstream and downstream of this population are polarised towards a more primitive, HSC/MPP-like, quiescent state on one side, and more downstream differentiated cells on the other. Surviving LSCs acquire a transcriptional profile consistent with enhanced long-term repopulating activity and stemness, and human mitochondrial DNA mutation and murine barcode analyses reveal that all cell clones surviving IFNγ display this fate bifurcation, contain highly functional LSCs and are therefore poised to remain long-term. Multigen functional analysis coupled with serial PDX assay confirms that IFNγ inhibits AML growth only temporarily, and in vivo murine models demonstrate the relevance of this phenomenon for AML growth. AML molecular and functional heterogeneity is a known challenge in defining universal behavior within this disease including the response to IFNγ challenge, and it is noteworthy that the pattern of primitive LSC enrichment coupled with ineffective progenitor proliferation was evident across AML samples with diverse cytogenetics. Future larger scale studies will be required to confirm the universality of these findings, and this survival mechanism is likely to contribute to the substantial proportion of AML relapse cases arising without further driver-mutations^53^.

Several treatments routinely used for AML generate inflammation. Graft versus leukaemia responses make HSC transplant the current most effective therapy, immunotherapy approaches by definition harness inflammation, and chemotherapy itself has also been shown to drive inflammation including IFNγ upregulation^69^. LSCs surviving IFNγ enter a deeper quiescent state both transcriptionally and functionally and this is likely to render them refractory to chemotherapy^12,54^. Indeed we have shown that they are enriched in signatures of LSCs resistant to the standard of care cytarabine therapy^12^. The IFNγ driven leukaemia stemness induction we identify is consistent with relapse AML being enriched in more primitive cells^54^. Our data suggest that chemotherapy efficacy may be affected when administered in combination or shortly following any interventions that induce inflammation. Our refined understanding will provide a workable framework to guide and instruct the design of better therapeutic schemes where chemotherapy and INFγ can synergise instead of antagonise together.

## Materials and Methods

### Primary AML samples and culture

Primary AML human samples were obtained and frozen at diagnosis, with informed consent at Hammersmith Hospital (London, UK) and provided from the Imperial College Healthcare Tissue and Biobank (ICHTB) with ethical approval by Wales REC3 (22/WA/0214), or at University College Hospital and provided from the UCLH Biobank with ethical approval by REC20/YH/0088. Details of patient samples are provided in Table 1. AML mononuclear cells were isolated by centrifugation using Ficoll-Paque (GE Healthcare Life Sciences). Cells were thawed and cultured on a monolayer of MS5 stromal cells in H51 Myelocult media supplemented with recombinant human 3GT cytokines (20ng/ml IL-3, 20ng/ml G-CSF, 20ng/ml TPO) and 50ng/ml recombinant human IFNγ or PBS control and kept in culture for 3 days at 37oC, 5% CO2. Cells were then harvested for FACS analysis, or to be used in the MultiGen assay (section below), or to sort CD34+ AML cells for snRNAseq&ATAC sequencing (section below). In some experiments, cells were thawed, FACS stained and FACS sorted for LSC-enriched AML cells (CD3^−^, CD19^−^, CD45^dim^, CD33^dim^, CD38^−^, CD34^+^ or CD3^−^, CD19^−^, CD45^dim^, CD33^dim^, CD38^+/−^, CD34^+^), to set up *in vitro* cultures in Stem Span medium (StemCell Technologies). In all experiments, cells were plated at 200,000 cells/ml. All cytokines were from Peprotech.

### Flow Cytometry and cell sorting

For the immunophenotypic analysis of T cells and AML cells, the following fluorochrome-conjugated primary antibodies specific to mouse were used: CD3e (145-2C11 or 17A2), CD8a (53-6.7), CD44 (IM7), PD1 (29F.1A12 or J43), PDL1 (M1H5 or 10F.9G2), c-Kit (2B8), CD11b (M1/70), Ly6C (HK1.4), CX3CR1 (SA011F11), Ki67 (11F6 or B56). Antibodies were purchased from BD Bioscience, Biolegend and eBioscience. Leukaemic cells were identified by the expression of tomato or YFP fluorescent proteins, as specified in individual experiments.

For the immunophenotypic analysis of human AML cells, the following fluorochrome-conjugated primary antibodies specific to human were used: CD3e (UCHT1), CD19 (4G7), CD33 (WM53 or HIM3-4), CD45 (HI30), CD38 (HIT2), CD34 (8G12), CD14 (clone), CD16 (clone), CD15 (clone) HLA-DR-DP-DQ (Tu39) and PDL1 (29E.2A3) antibodies. Live and dead cells were distinguished using 7AAD (Biolegend or BD Biosciences), or fixable viability dyes 780 or BV510 (BD Biosciences). Calibrite beads (BD Biosciences) were used to determine absolute cell counts in specified experiments. Cells were analysed with a BD Symphony or LSR Fortessa, and sort purified using a FACSAria or Fusion (BD Biosciences). Data were analysed with FlowJo (TreeStar).

### scRNA sequencing analysis

#### Human Dataset

##### Single-nuclei RNA-seq data processing

4 primary AML samples (for patient details see table 1) were cultured in H51 myelocult supplemented with human recombinant 3GT cytokines and 50ng/ml IFNγ or PBS control, on a monolayer of MS5 stromal cells and 3 days later, cells were harvested, FACS stained and sorted for live, CD45dim, CD33positive/dim, CD14-, CD16-, CD15-, CD38+/− and CD34+ cells. Single nuclei 10x Multiome sequencing data (snRNA-seq + snATAC-seq) were processed using Cell Ranger ARC (v.2.0.1, 10x Genomics) with default settings. Reads were aligned to the prebuilt Human reference (refdata-cellranger-arc-GRCh38-2020-A-2.0.0). Bulk RNA-seq data were processed using the nf-core RNAseq pipeline (revision 3.14.0, https://github.com/nf-core/rnaseq). Alignment was performed using STAR^55^ against the Human genome build GRCh38 release-98 (consistent with the reference used in Cell Ranger ARC), and quantification was conducted using RSEM^56^.

#### Demultiplexing of 10x Multiome Data

To demultiplex 10x multiome samples and accurately assign nuclei back to their respective patient donor, we used genotype information from matched bulk RNA-seq data for each donor. Specifically, RNA-seq bam files were first genotyped using Cellsnp-lite v.1.2.3^57^ to identify individual-specific SNPs, which served as a reference for demultiplexing. Cellsnp-lite was next used to pile up these variants on the polled snRNA-seq bam files generated by cellranger. Finally, these pileups, together with the donor-specific genotypes, were used to demultiplex samples and assign each nucleus to a donor using Vireo v.0.5.8^58^.

#### Demultiplexing of 10x Multiome Data using mitochondrial mutations

mgatk v0.6.1 (Lareau, C. (2020). *mgatk: mitochondrial variant calling tool*. GitHub repository. Retrieved from https://github.com/caleblareau/mgatk) on the scATAC-seq data was ran separately for each condition to generate mitochondrial allele count tables, which were then merged for downstream demultiplexing analysis. Cells with a mean mitochondrial read depth below 5 were excluded to ensure reliable variant detection. The filtered allele count table and cell list were used as input to scMitomut v1.4.0 (Sun W (*2025). scMitoMut: Single-cell Mitochondrial Mutation Analysis Tool.* doi:10.18129/B9.bioc.scMitoMut*, R package version 1.4.0,* https://bioconductor.org/packages/scMitoMut.), a tool designed to detect mitochondrial mutations across single cells. The following parameters were set: min_cell = 11, requiring each variant to be present in at least 11 cells, and p_threshold = 0.01 to define the significance threshold for mutation detection. This process produced two matrices: an allelic frequency matrix and a q-value matrix. In both matrices, rows correspond to mitochondrial loci and columns to individual cells. Each entry in the allelic frequency matrix represents the proportion of wild-type reads at a given locus in a given cell, while each entry in the q-value matrix reflects the probability that the corresponding locus is not mutated in that cell.

To further refine donor demultiplexing, donor sex was incorporated as an additional layer of deconvolution. The sex of each cell was determined using gene expression profiles obtained from the RNA data. Specifically, cells were labeled as: Female if expressing the XIST gene; Male if expressing any of the male-specific genes RPS4Y1, DDX3Y, UTY, or KDM5D; Doublet if both XIST and at least one male-specific gene were expressed; Unknown if none of the sex marker genes were detected. Each cell was then classified as consistent, inconsistent, doublet, or unknown by comparing the inferred sex with the donor’s annotated sex. A cell was considered **consistent** if the inferred and annotated sexes matched, and **inconsistent** if they did not. Only consistent cells were retained for subsequent mtDNA analysis. Finally, to verify the SNP-based deconvolution, the retained cells were clustered using the q-value matrix. Hierarchical clustering was performed with complete linkage and Euclidean distance as the metric using the pheatmap package (Kolde, R. (2019). *pheatmap: Pretty Heatmaps*. R package version 1.0.12. Retrieved from https://cran.r-project.org/package=pheatmap). Clusters were annotated with donor identities obtained from SNP-based deconvolution to evaluate concordance between the two approaches (Extended data Fig.8a).

#### Single-nuclei RNA-seq data processing

Raw sequencing reads were aligned to GRCh38 using cellranger-arc with the depth normalisation method to equalize for the mean number of reads that are confidently mapped to the transcriptome per cell for each gene expression library and lane, to correct for read depth coverage between the different lanes samples were sequenced on.

Counts from gene expression were extracted from the multiome datasets to create a Seurat object. Mitochondrial and ribosomal transcript percentages were calculated. Data were log-normalized using the NormalizeData function and scaled using the ScaleData functions from the Seurat package. Only correct cells, as described in the section “Demultiplexing of 10x Multiome Data using mitochondrial mutations” above for which patient were confidently identified and were only singlets, as described in the same section were retained for the analysis.

#### Reference map

Cells that were successfully identified as “correct” as previously described in the section Demultiplexing of 10x Multiome Data using mitochondrial mutations were mapped to the reference map^54^ using Symphony and Patient ID as batch correction.

Quality controls were run based on mapping error score and cells with mapping errors>= to 2.5 MADs above median were excluded. Mapping error was calculated using the calculate_MappingError function.

This percentage remained small for every patient sample. Allocation of leukemic cells to their closest population in the normal hematopoietic hierarchy was determined using KNN classification and the predict_CellTypes function. Cells were plotted on the reference map using the ggplot package or Seurat packages.

Pseudotime score corresponding to the level of differentiation from normal HSC to any mature cells in the reference map was determined using the predict_Pseudotime function from the BoneMarrowMap package. As such, this score enables to determine how far from the normal hematopoietic stem cell population the cell of interest is along the hemopoietic differentiation trajectory. For some analyses, cells were grouped in 3 states. HSC-like states which included: HSC/MPP, cells grouped in progenitor-like states included: cycling progenitor, early erythroid, early GMP, early Lymphoid, Eosinophil-Basophil-Mast cell Precursor, late GMP, LMPP, Megakaryocyte Precursor, MEP and differentiated-like states included: B cell, late erythroid, monocyte, naïve T cell, pDC, pre-B cell, Pro-B cell, pro-monocyte, CD4 Memory T cell, CD8 memory T cell, cDCs.

UMAP were visualized using ggplot function or the built in functions in the Seurat package. Predicted pseudotime density was plotted using the ggdensity package.

AUC scores were calculated for the stated signatures using the AUC scoring method (https://bioconductor.org/packages/release/bioc/html/AUCell.html), *to identify cells that are actively expressing genes* within each of the gene list depicted in the results section.

Cell cycle assessment

Cell cycle status was assessed using the CellCycleScoring function from the Seurat package. AUC score was further calculated using the R package AUCell (https://bioconductor.org/packages/release/bioc/html/AUCell.html), for the cell cycle signatures stated in the figures.

#### List of signatures used

S Score and G2M scores were calculated using the Seurat package with the function CellCycleScoring and default parameters.

B1/2/3 signatures were generated comparing all cells part of Branches 1, 2 and 3 (Stem cell branches) to cells part of the other branches (non-stem cell branches) from the multiome dataset generated from the OCI-AML22 CD34+CD38-fraction in Boutzen et al, 2024, Leukemia^12^. Genes with pvalues<0.05 were selected. B1/2/3 up represents the list of genes upregulated (log2 fold change >0) while B1/2/3 down signature represents the list of genes downregulated (log2 fold change <0).

B1/2 signatures were generated comparing all cells part of Branches 1, 2 from Boutzen et al, 2024^12^. These corresponds to Stem cell branches that are associated with the deepest level of quiescence, the slowest to repopulate mice and the highest level of resistance to cytarabine and whose signature is the most associated with poor prognosis) to cells part of the other branches (non-stem cell branches) from the multiome dataset generated from the OCI-AML22 CD34^+^CD38^−^ fraction in Boutzen et al, 2024^12^. Genes with pvalues<0.05 were selected. B1/2 up represents the list of genes upregulated (log2 fold change >0) while B1/2 down signature represents the list of genes downregulated (log2 fold change <0).

The Hallmark IFNγ response signature was extracted from the Molecular Signatures Database (MSigDB)^59^.

#### Mitochondrial mutation encoding and clustering analysis

To identify putative clones using mitochondrial mutations, we constructed a binary mutation matrix and applied a clustering analysis.

Starting from the q-value matrix aggregated across donors, we separated it into donor-specific matrices. Each matrix was then trinarized based on q-values: loci with q-value < 0.05 were coded as mutated (1), q-values > 0.95 as non-mutated (0), and intermediate q-values as uncertain (NA). For each locus, we counted the number of cells in the mutated (1) and non-mutated (0) states, considering only cells with observed values and excluding missing data. When the mutated state was present in fewer cells than the non-mutated state, it was kept encoded as 1. When the non-mutated state was present in fewer cells, labels were swapped so that the less frequent cell state was always encoded as 1, corresponding to the mutated state. Then, uncertain entries (NA) were recoded as 0, resulting in a binary matrix where 1 indicates mutated cells and 0 indicates non-mutated or uncertain cells. Loci with fewer than 10 mutated cells were removed, as well as cells that were never mutated at any locus. Clustering was performed using bootstrap resampling using the pvclust R package version 2.2 (1) (nboot = 500), with binary distance (Jaccard) and average linkage. Clusters were retained if they showed strong bootstrap support (AU p-value ≥ 0.95) and included more than 10 cells in total, with at least 2 cells per condition.

#### Murine dataset

Processed data from murine MLL-AF9-Kras*^G12D^* AML cells transplanted into NSG and C57Bl/6 recipients and harvested from the spleens of fully infiltrated animals with matched SPLINTR barcodes^44^ were kindly provided by Dane Vassalides and Mark Dawson. Splenic AML cell scRNAseq datasets from NSG (n=2) and C57Bl/6 (n=2) mice were integrated according to standard Seurat workflow. Cell passing quality control thresholds for number of reads and mitochdonrial transcript proportion were retained for downstream analysis. SCTransform normalisaiton was performed on each sample and 3,000 features were selected for integration. Clonal abundance was quantified based on the proportion of SPLINTR barcoded clones, identified by barcode expression, within each scRNAseq dataset. Transcriptional signature scores were calculated using Vision v.2.1.0^60^ using the LSC signature identified in MLL-AF9 cells^61^ and the Hallmark Interferon Gamma Response Mouse signature^59^. To obtain monocyte signature scores, SingleR v1.8.1 and CellDex v1.4.0^62^ packages were used to calculate cell-type specific scores based on ImmGen Mouse database^63^ from which monocyte-specific signature scores were extracted.

#### MultiGen assay

The assay was performed as previously described^46,47^. Briefly, Five primary AML samples (for patient details see table 1) were cultured on a monolayer of MS5 stromal cells, in H51 myelocult medium (Stem Cell Technologies), supplemented with recombinant human 3GT cytokines (20 ng/ml IL-3, 20 ng/ml G-CSF, 20 ng/ml TPO) and 50 ng/ml recombinant human IFNγ or PBS control, and kept in culture for 3 days at 37°C, 5% CO2. PBS/IFNγ pre-treated primary AML cells were labelled with one of four cell trace label combinations (2.5μΜ CTVhigh only, 2.5μΜ CFSEhigh only, 2.5μΜ CTVhigh and 1.25μM CFSElo together, 1.25μM CTVlo and 2.5μΜ CFSEhigh together) and FACS stained for CD34 and CD38. CellTrace labels were purchased from Thermofisher. Four CD34^+^ AML cells, each one with a distinct label, were then FACS sort-plated per well and a bulk population of CD34^+^ AML cells including all cell trace combinations was plated in a separate well (1000-3000 cells per cell trace combination and condition). Everything was cultured for 72 hours in StemSpan Serum Free (SFEM) medium (Stem Cell Technologies) supplemented with recombinant human SCF 50 ng/mL, FLT3l 50 ng/mL, TPO 20 ng/mL, IL-3 10 ng/mL, IL-6 10 ng/mL, G-CSF 10 ng/mL and 50ng/ml recombinant human IFNγ or PBS control, and kept in culture for 3 days at 37°C, 5% CO2. At this point cells in each well were stained with CD34 and CD38 antibodies and their phenotype and cell trace label dilution was assessed by FACS analysis, using the ZE5 flow cytometer (BioRad) which enables the analysis of all contents of each well. The data obtained from each well plated with 4 cells were mapped on the data from the bulk cultures. In the single cell data, the cell trace label serves to identify cell families originating from each ancestor CD34^+^ cell in each well, while the dilution of the label identifies the number of divisions undergone by individual cells within each clonal family (generation) and the CD34 phenotype reports on the putative stem/differentiation state of each cell recovered. The MDN was calculated based on the Precursor Cohort method, as previously described: shortly, it removes the effect of cell division on the final cell numbers, converting the number of cells for a given generation to a precursor cohort number^84^.

### Mice

Animal work was performed in accordance with the animal ethics committee (AWERB) at Imperial College London and the Francis Crick Institute, UK and following UK Home Office regulations (ASPA, 1986). NOD-scid IL2Rgamma^null^ (NSG), and C57Bl/6 mice were bred and housed at the Francis Crick Institute, in individually ventilated cages. Aged-matched, female mice >8 weeks old were used for all experiments.

#### NP-MLL-AF9 murine AML model

Primary AML cells were generated from granulomonocytic progenitors (GMPs) from mT/mG or PU.1-YFP mice (C57Bl/6 background) using MLL-AF9-IRES-GFP retroviral particles as described^13^. BM cells and splenocytes were harvested from primary mice, when fully infiltrated with leukemia (BM AML infiltration>90%) and were cryopreserved as individual batches of primary AML cells^13^. To generate AML cells expressing the NP-model antigen, PlatE adherent packaging cells were transfected using Calcium phosphate precipitation (Invitrogen) with the pMP71 NP-ires-GFP construct (kind gift from H.J.Stauss). Viral supernatant was harvested from the transfected cells and used to transduce primary BM AML cells by spinfection on retronectin-coated (Takara-Bio Inc) plates. The transduced NP-AML cells were expanded *in vitro* and GFP bright cells, which expressed the model antigen, were FACS purified and injected intravenously for *in vivo* expansion in NSG mice. Spleens and BM were harvested from fully infiltrated animals and single cell suspensions from each mouse were frozen as individual batches of NP-AML cells. Primary AML cells were thawed, washed and resuspended in PBS and up to 100,000 cells were injected into secondary syngeneic C57Bl/6 recipients or immunodeficient NSG mice, as described in individual experiments. Experimental mice receiving NP-MLL-AF9 AML (NP-AML) cells were never conditioned. Leukaemic animals were sacrificed when AML infiltration in the blood was more than 20% or when they exhibited the first clinical signs of high AML infiltration.

#### Tissue harvest and single cell suspension preparation

Blood was sampled either by venipuncture of the mouse tail, or through cardiac bleeds following CO2 administration and collected into heparin-coated tubes to prevent clotting. To prepare single cell suspensions of splenocytes, freshly harvested spleens were mashed with the plunger of an 1ml syringe, in RPMI supplemented with 10% FBS and passed through a 70-μm cell strainer. To prepare single cell suspensions of BM cells, femurs, tibias and pelvic bones were crushed with a pestle and mortar in RPMI medium supplemented with 10% FBS and passed through a a 70-μm cell strainer. Red blood cells were removed by isotonic lysis with ammonium chloride. Cells were resuspended in FACS buffer (PBS, 2% FCS) for counting, immunolabelling and FACS analysis. In some experiments, BM, splenocytes and blood from individual leukaemic animals were cryopreserved in freezing media (10% DMSO and 90% FBS) and thawed for further analyses at later points.

#### Murine AML cell culture

AML cells were plated at 0.25×10^5^/ml in RPMI medium, supplemented with 10% FBS, 1% penicillin/streptomycin, 1% L-glutamine and 10ng/ml recombinant murine SCF, 6ng/ml recombinant murine IL-3 and 10ng/ml recombinant human IL-6. When specified, 10ng/ml recombinant murine IFNγ was added to the cultures. All cytokines were purchased from Peprotech. For co-culture experiments, 100 000 NP-AML AML cells harvested from the spleens of fully infiltrated leukaemic mice were plated in the above media, together with splenocytes from F5 or OTI TCR transgenic mice at a1:2 ratio, in 96-well u-bottom plates, in 200μl/well for 4 days. In some experiments, AML cells were cultured in the above media with the addition of conditioned media (1:1 volume) from F5:NP-AML cocultures, or OTI:NP-AML cocultures, in the presence or absence of an IFNγ blocking antibody (R4-6A2 clone, NA/LE, BD Biosciences). Cells were incubated at 37°C, 5% CO2.

#### Colony Forming Unit Assays

AML cells were stained with c-Kit and PDL1 antibodies and FACS sorted based on their phenotype as specified, directly into Methocult^TM^ (M3434) and plated into 6-well plates. In single cell colony forming assays, c-Kit^+^ AML cells were FACS sorted directly into 96-well plates containing Methocult^TM^ (M3434), so that 1 cell/well was plated. Cultures were incubated at 37°C and quantified on days 6 to 9, as specified in individual experiments. To plate single, c-Kit+ AML cells into secondary colonies, 3-5 primary colonies per mouse were harvested, FACS stained and 12 c-Kit+ cells from each primary colony were plated at 1 cell per well in a 96-well plate of methocult. In total 300 cells plated from NSG leukaemic mice and 300 cells plated from C57Bl/6 recipients were plated into secondary colonies and counted 8 days after plating.

### Statistical analysis

For MultiGen assay data, statistical significance on the data presented in Figure 3e, as previously described^46,47^, In brief, a permutation-based test to challenge the null hypothesis that the distribution of the progeny recovered from the multiGen assay in different generations was independent of IFNγ exposure was performed. For all other data, statistical analysis was performed using GraphPad Prism (GraphPad Software). For all data, differences were considered significant whenever p<0.05. * p <0.05, ** p< 0.01, *** p < 0.001, ****p<0.0001. Number of animals and samples, and statistical tests used are specified in Figure legends.

---

**Supplementary Figure 1.**
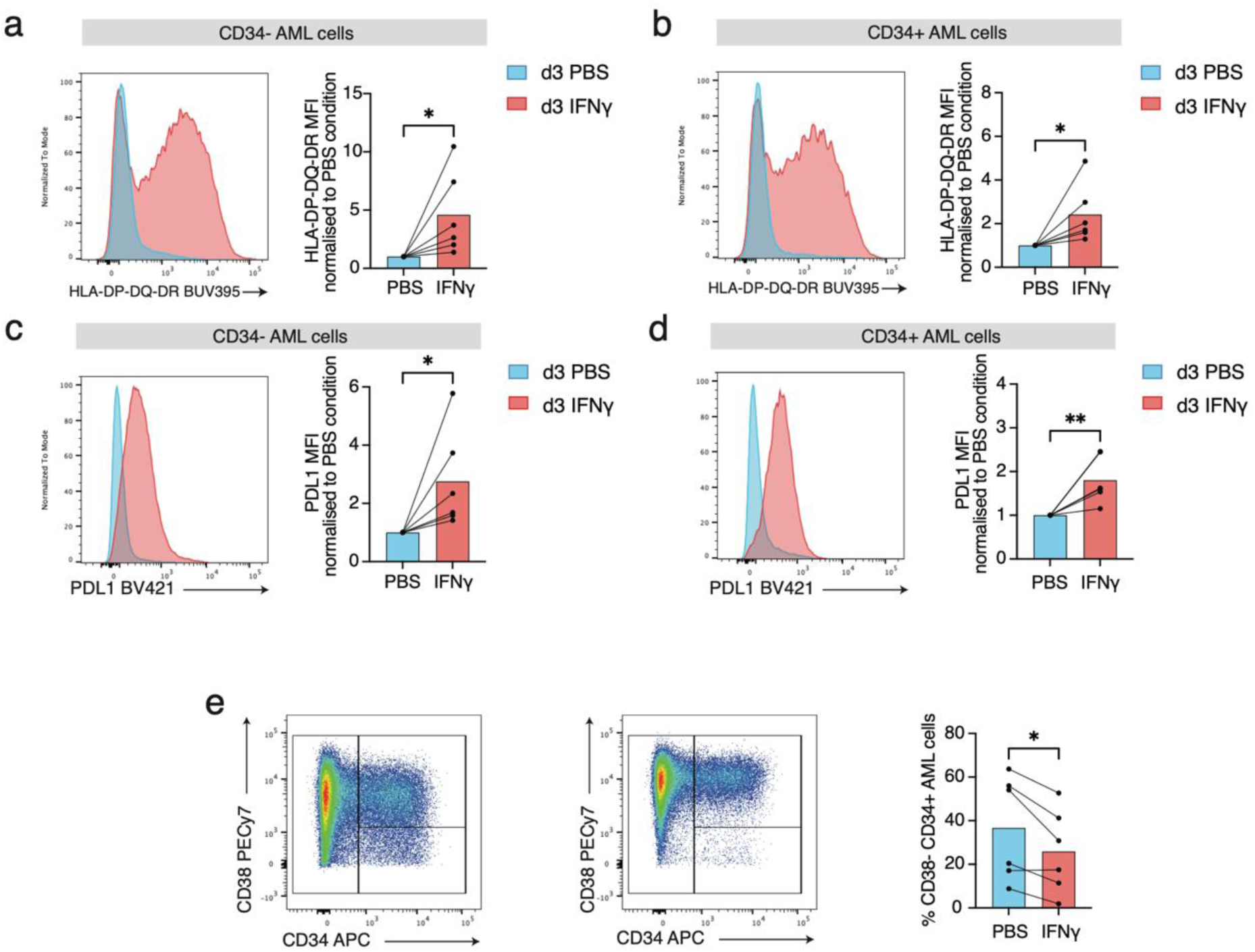
**a-b,** Representative FACS histogram overlays showing HLA-DP-DQ-DR expression on CD34^−^ (a) and CD34+ (b) cells from primary AML cells cultured in the presence of IFNγ (red) or PBS control (blue) for 3 days. Adjacent bar graphs summarise fold-change in HLA-DP-DQ-DR expression following IFNγ treatment. **c-d,** Representative FACS histogram overlays showing PDL1 expression on CD34^−^ (c) and CD34+ (d) cells from primary AML cells cultured in the presence of IFNγ or PBS control for 3 days. Adjacent bar graph summarises fold-change in PDL1 expression following IFNγ treatment. **e,** Representative FACS plots showing CD34 and CD38 expression among primary AML cells cultured in the presence of PBS control (left) or IFNγ (right) for 3 days. Adjacent bar graph summarising the percentage of CD38^−^ CD34^+^ cells among AML cells in the two conditions. n=6 patient samples. Each dot represents a patient sample. Two-tailed paired t-test performed for statistical significance.

**Supplementary Figure 2.**
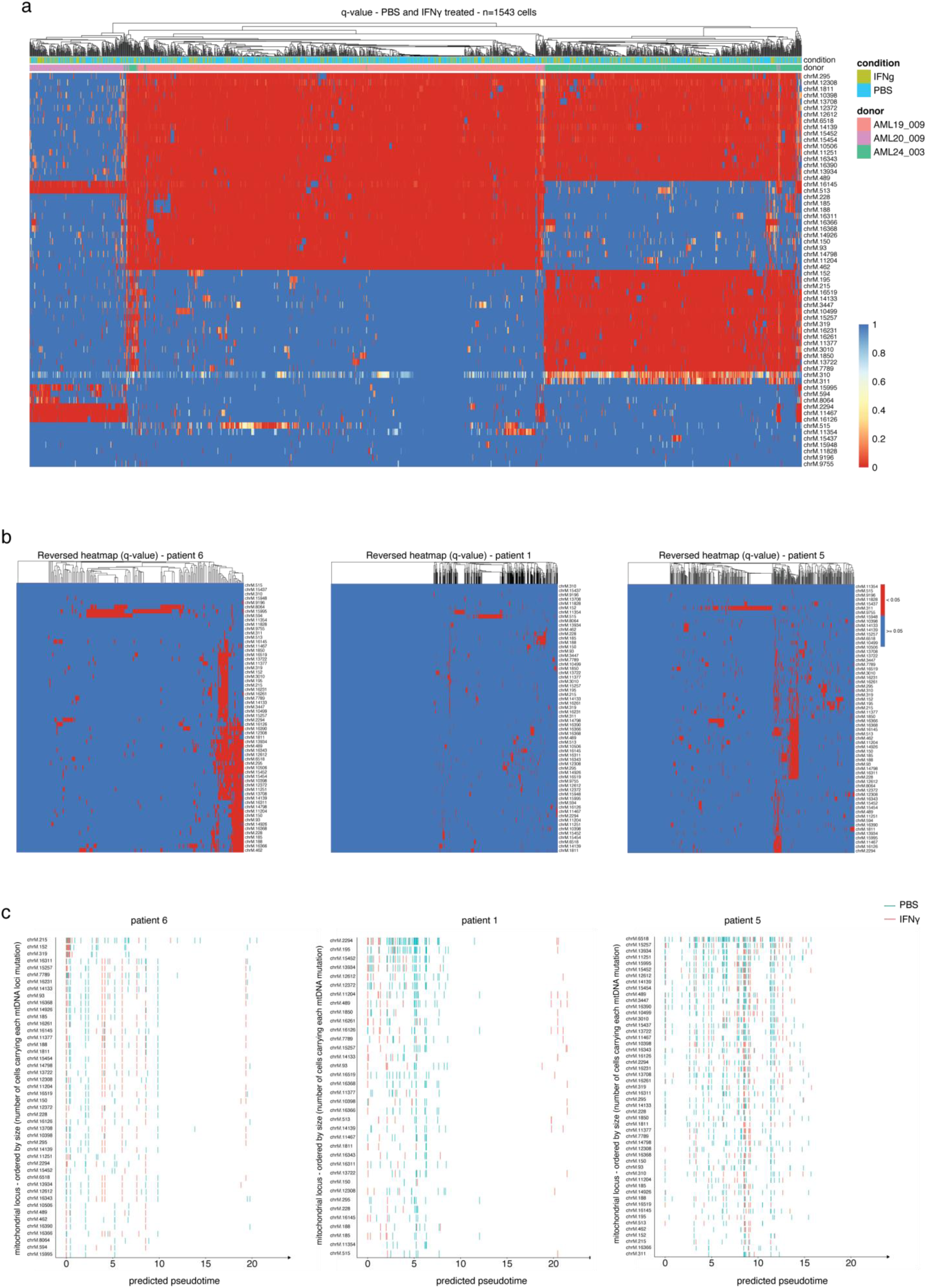
**a,** Heatmap showing q-values (probability of no mutation) for each mitochondrial locus in each cell, annotated by patient and condition. Columns represent cells and rows represent mitochondrial loci. Lower q-values indicate a higher probability of a mutation at that locus. Only cells with at least one mitochondrial mutation and with concordant chromosomal and donor sex are shown. **b,** Heatmap showing the binarized mutation status for each mitochondrial locus in each cell, shown for each patient. Columns represent cells and rows represent mitochondrial loci. Loci with q-value < 0.05 are shown in red (mutated), while loci with q-value ≥ 0.05 are shown in blue (non-mutated or uncertain). **c,** Distribution of mitochondrial clones along predicted pseudotime (defined in Extended Data Fig. 7c). Each row represents a mitochondrial locus, x-axis represents pseudo-time, and bar colors indicate individual cells carrying the specified mitochondrial mutation in each condition (PBS in blue, IFN-γ in red). Only loci with at least 10 mutated cells are shown, and loci are ordered by clone size.

**Supplementary Figure 3.**
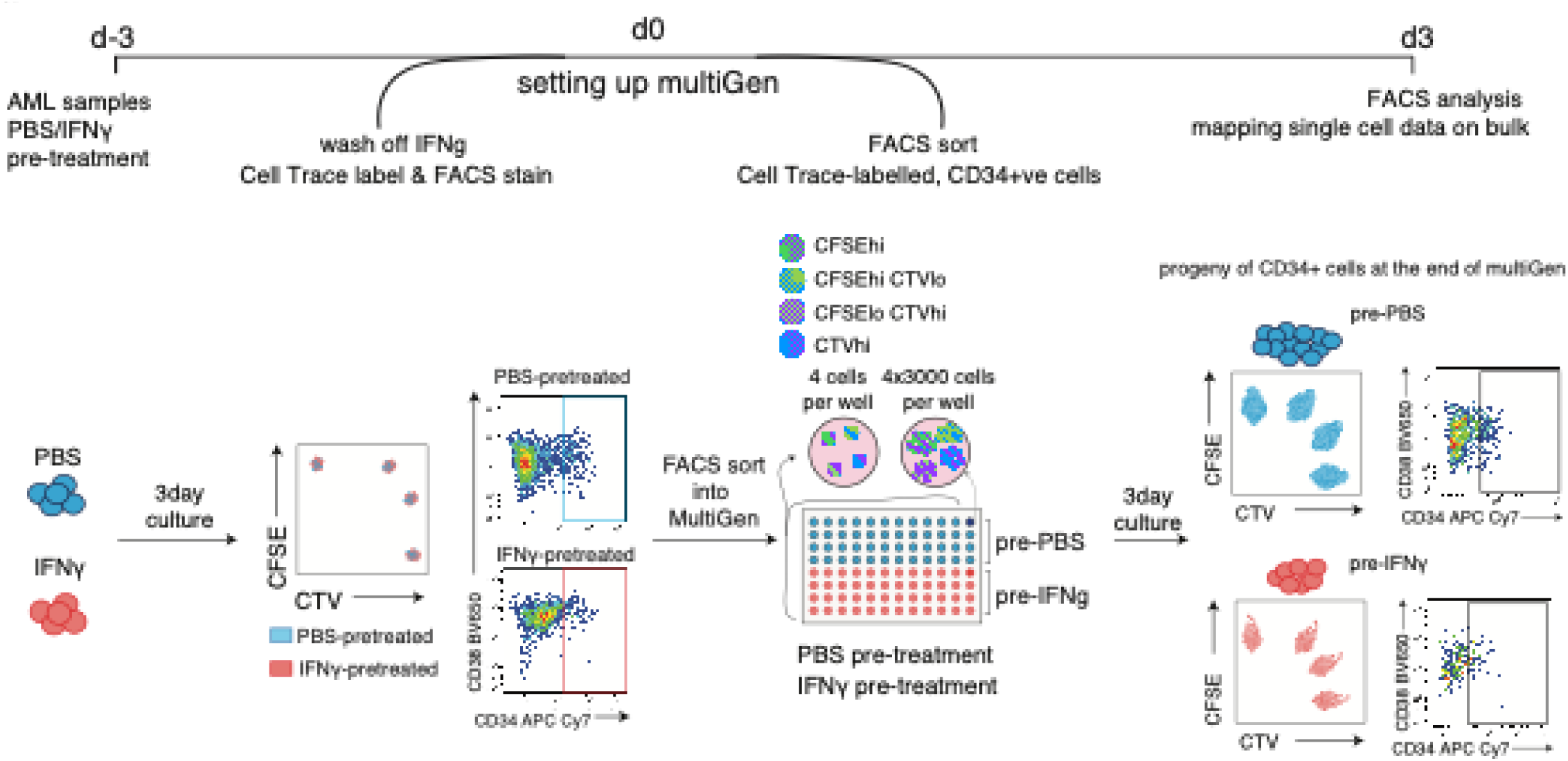
Schematic diagram describing experimental set up and multiGen assay. Primary AML samples were pre-treated with PBS or 50ng/ml rhIFNγ for 3 days (d-3 to d0). On day 0, the cells were harvested, IFNγ was washed off and AML cells were cell trace labelled with one of four cell trace label combinations (CTVhigh only, CFSEhigh only, CTVhigh and CFSElo together, CTVlo and CFSEhigh together). Four CD34^+^ AML cells (CD33dim, CD45dim, CD34+), each one with a distinct label, were then FACS sort-plated per well and a bulk population of CD34^+^ AML cells including all cell trace combinations was plated in a separate well. Everything was cultured for 72 hours and at this point all resulting cells in each well were analysed using flow cytometry. The data obtained from each well plated with 4 cells were mapped on the data from the bulk cultures. In the single cell data, the cell trace label serves to identify cell families originating from each ancestor CD34^+^ cell in each well, while the dilution of the label identifies the number of divisions undergone by individual cells within each clonal family (generation) and the CD34 phenotype reports on the putative stem/differentiation state of each cell recovered.

**Supplementary Figure 4.**
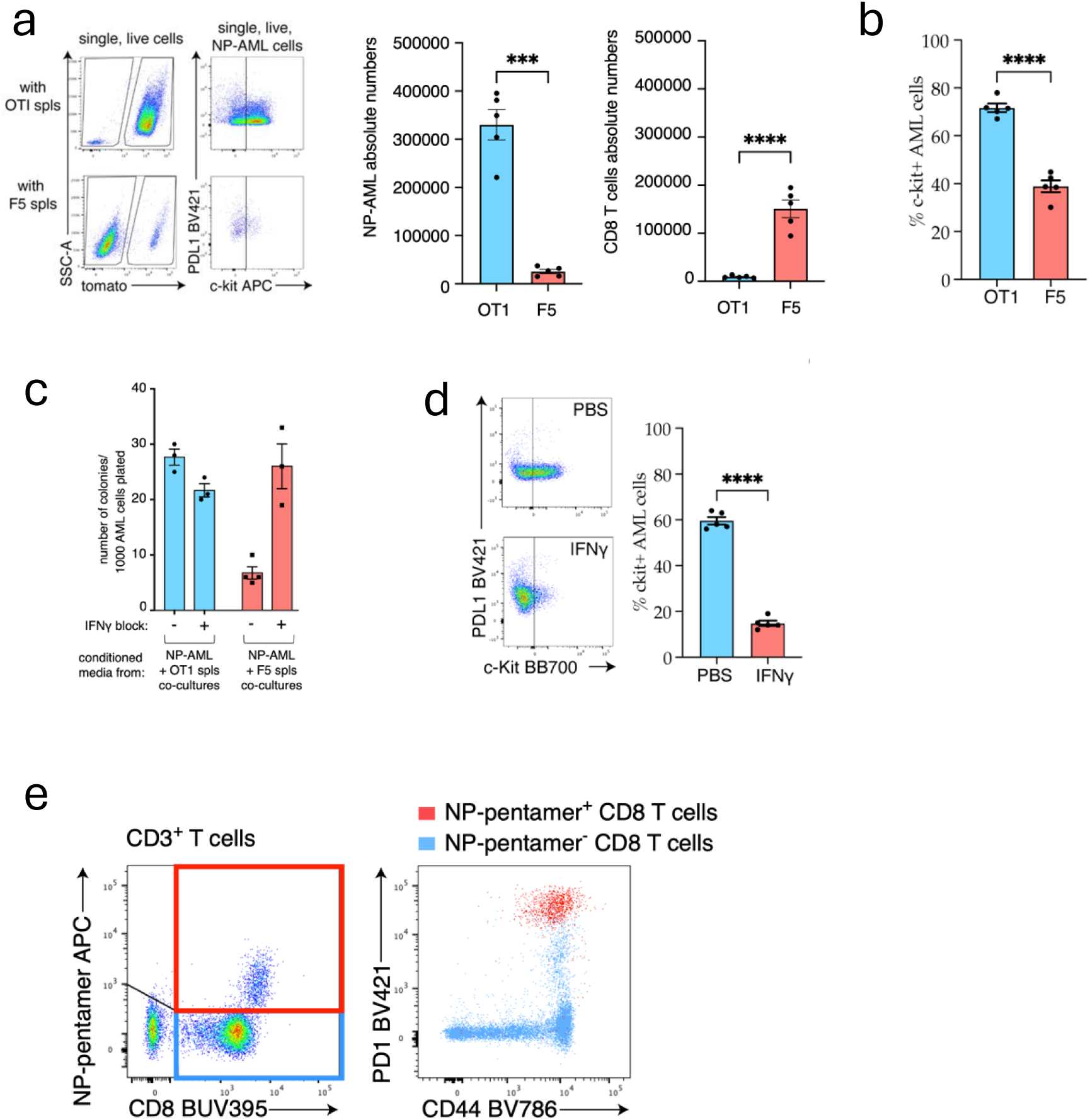
**a,** Representative FACS plots of NP-AML cell: OTI/ F5 TCR tg splenocyte co-cultures, showing tomato+ AML cells (left plots) and PDL1 and c-Kit expression (right plots). Bar graphs indicate (left) absolute number of NP-AML cells and (right) absolute number of CD8 T cells in the co-cultures. **b,** percentage of c-Kit+ NP-AML cells in each co-culture. a-b, N=5 NP-AML cell samples, harvested from 5 different NSG leukaemic mice. Dots are the average of technical triplicates of OT1/F5 splenocyte co-cultures and each dot represents co-cultures of NP-AML cells from individual NSG donors. Error bars: mean± SEM. Paired t-tests used for statistical significance. **c,** Colony forming assays NP-AML cells cultured for 3 days in conditioned media (1:1 volume) from F5/OTI:NP-AML cocultures, in the presence or absence of IFNγ blocking antibody. N=3 co-cultures/group. Error bars: mean ± SEM. **d,** Representative FACS plots of NP-AML cells harvested from NSG recipients and treated with IFNγ or PBS control for 18hrs. Bar graph summarising the percentage of c-kit+ AML cells in each condition, each dot represents NP-AML cells harvested from a different leukaemic mouse, error bars: mean ± SEM, paired t-tests were performed for statistical significance. **e,** Representative FACS plots of T cells in the blood of C57Bl/6 animals fully infiltrated with NP-AML. Left: NP-pentamer and CD8 staining on live, single, CD3^+^ cells (CD8 T cells). Right: overlay of NP-specific T cells (live, single, tomato^−^, CD3^+^, CD8^+^, NP-pentamer^+^ cells – red gate on the left FACS panel) and remaining, CD8 T cells (live, single, tomato^−^, CD3^+^, CD8^+^, NP-pentamer^−^ cells – blue gate on the left FACS panel), showing PD1 and CD44 expression in each population.

